# Dynamin binding protein is required for *Xenopus laevis* kidney development

**DOI:** 10.1101/414458

**Authors:** Bridget D. DeLay, Tanya A. Baldwin, Rachel K. Miller

**Affiliations:** Department of Pediatrics, Pediatric Research Center, University of Texas Health Science Center McGovern Medical School, Houston, TX, USA; Department of Integrative Biology and Pharmacology, McGovern Medical School, University of Texas Health Science Center, Houston, TX 77030, USA; Program in Biochemistry and Cell Biology, University of Texas MD Anderson Cancer Center University of Texas Health Science Center Graduate School of Biomedical Sciences, Houston, TX, USA; Program in Genetics and Epigenetics, University of Texas MD Anderson Cancer Center University of Texas Health Science Center Graduate School of Biomedical Sciences, Houston, TX, USA; Department of Genetics, University of Texas MD Anderson Cancer Center, Houston, TX, USA

**Keywords:** Dnmbp, Tuba, *Xenopus*, nephrogenesis, pronephros, CRISPR

## Abstract

The adult human kidney contains over one million nephrons, with each nephron consisting of a tube containing segments that have specialized functions in nutrient and water absorption and waste excretion. The embryonic kidney of *Xenopus laevis* consists of a single functional nephron composed of regions that are analogous to those found in the human nephron, making it a simple model for the study of nephrogenesis. The exocyst complex, which traffics proteins to the cell membrane in vesicles via CDC42, is essential for normal kidney development. Here, we show that the CDC42-GEF, dynamin binding protein (Dnmbp/Tuba), is essential for nephrogenesis in *Xenopus*. *dnmbp* is expressed in *Xenopus* embryo kidneys during development, and knockdown of Dnmbp using two separate morpholino antisense oligonucleotides results in reduced expression of late pronephric markers, whereas the expression of early markers of nephrogenesis remains unchanged. A greater reduction in expression of markers of differentiated distal and connecting tubules was seen in comparison to proximal tubule markers, indicating that Dnmbp reduction may have a greater impact on distal and connecting tubule differentiation. *dnmbp* knockout using CRISPR results in a similar reduction of late markers of pronephric tubulogenesis. Overexpression of *dnmbp* in the kidney also resulted in disrupted pronephric tubules, suggesting that *dnmbp* levels in the developing kidney are tightly regulated, with either increased or decreased levels leading to developmental defects. Together, these data suggest that Dnmbp is required for nephrogenesis.

## INTRODUCTION

Kidney development is conserved in amphibians and mammals, making *Xenopus* embryos a good model for studying nephrogenesis. Mammalian kidney development proceeds through three stages: the pronephros, mesonephros, and metanephros (Vize et al., 1997). Similarly, amphibian embryos have a pronephros, and adults have a metanephros (Vize et al., 1995; Vize et al., 1997). The basic unit of filtration for all kidney forms is the nephron, with the same signaling cascades and inductive events leading to nephrogenesis in mammals and amphibians (Brandli, 1999; Hensey et al., 2002). The *Xenopus* pronephros consists of a single, large, functional nephron (Brennan et al., 1998; Carroll et al., 1999), making it a simple model for studying vertebrate nephron development. Additionally, the *Xenopus* tadpole epidermis is transparent and the kidney is located just under the epidermis, allowing visualization of the pronephros without dissection (Carroll et al., 1999). It is also possible to easily modulate gene expression in *Xenopus* embryos through overexpression, knockdown and knockout experiments via microinjection of RNA constructs, antisense morpholino oligonucleotides (MOs) and CRISPR constructs (Corkins et al., 2018; DeLay et al., 2018; Miller et al., 2011). The established cell fate maps of the early *Xenopus* embryo facilitate tissue-targeted modulation of gene expression by microinjection into the appropriate blastomere (DeLay et al., 2018; DeLay et al., 2016; Moody, 1987a; Moody, 1987b). Taken together, *Xenopus* is a powerful model for studying essential nephrogenesis genes.

One gene that plays an essential role in kidney development is *cdc42*. Cdc42 is a Rho family small GTPase that was first discovered in *Saccharomyces cerevisiae* (Johnson and Pringle, 1990). It plays a role in cell migration, polarity, differentiation and proliferation, as well as branching of blood vessels and regulation of actin dynamics (Lavina et al., 2018; Melendez et al., 2013; Mizukawa et al., 2017; Nguyen et al., 2017; Schulz et al., 2015). Cdc42 is a molecular switch that cycles between active (GTP-bound) and inactive (GDP-bound) states through its interaction with guanine exchange factors (GEFs) and GTPase activating proteins (GAPs) (Bishop and Hall, 2000; Schmidt and Hall, 2002). While GAPs increase the intrinsic GTPase activity of CDC42, GEFs exchange GDP bound to Cdc42 for GTP and assemble complexes between Cdc42, scaffold proteins and kinases (Cerione, 2004).

Loss of Cdc42 in the mouse ureteric bud leads to abnormal nephron tubulogenesis due to branching, polarity and cytoskeletal defects, while loss of Cdc42 in the mouse metanephric mesenchyme results in failure of the renal vesicle and S-shaped body to develop (Elias et al., 2015). Similarly, loss of Cdc42 in the distal tubules of mouse kidney leads to death within a few weeks of birth due to kidney failure, cyst development and a decrease in ciliogenesis within the kidney cysts (Choi et al., 2013). Knockdown of Cdc42 via MO in zebrafish leads to dilated kidney tubules, glomerulus defects and disorganized cilia within in kidney tubules (Choi et al., 2013).

Although Cdc42 localizes on the apical surface of the kidney tubule epithelium, it needs to be activated by a GEF in order to regulate tubulogenesis and ciliogenesis (Martin-Belmonte et al., 2007; Zuo et al., 2011). Dynamin binding protein (Dnmbp, Tuba) is a Cdc42-specific GEF that is known to be concentrated on the apical surface of kidney epithelial cells (Otani et al., 2006; Qin et al., 2010). Knockdown of Dnmbp in MDCK cells leads to a decrease in cilia, polarity defects and inhibition of tubulogenesis, similar to that seen when Cdc42 is knocked down (Baek et al., 2016; Zuo et al., 2011). Here, we demonstrate that knockdown, CRISPR knockout and overexpression of *dnmbp* lead to tubulogenesis defects in *Xenopus* pronephric kidneys, indicating that this protein is required for nephrogenesis.

## MATERIALS AND METHODS

### Embryos

Adult pigmented *X. laevis* were purchased from Nasco (LM00531MX). Eggs were obtained from female frogs, fertilized *in vitro* and the embryos reared as described previously (Sive et al., 2000). The Center for Laboratory Animal Medicine Animal Welfare Committee at the University of Texas Health Science Center at Houston, which serves as the Institutional Animal Care and Use Committee, approved this protocol (protocol #AWC-16-0111).

### Western blots

Embryos were collected at various stages (Nieuwkoop and Faber, 1994) for lysate creation. Protein lysates from 20 pooled embryos of the same stage were created as described previously (Kim et al., 2002), and one embryo equivalent was added per lane of an 8% SDS-PAGE polyacrylamide gel. Following transblotting of the protein onto a 0.45 µm PVDF membrane (Thermo Scientific), the blot was blocked for 3 hr in KPL block (SeraCare) at room temperature. After blocking, the membrane was incubated overnight at 4°C in 1:500 mouse anti-Dnmbp antibody (Abcam 88534) or 1:1000 rabbit anti-GAPDH antibody (Santa Cruz FL-335). Blots were rinsed with TBST and incubated in 1:5000 goat anti-mouse or goat anti-rabbit IgG horseradish peroxidase secondary antibody (BioRad, Hercules, CA) for 2 hr at room temperature. Blots were rinsed again in TBST and imaged using enhanced chemiluminescence (Pierce Supersignal West Pico) on a BioRad ChemiDoc XRS+.

### In situ hybridization

A DIG RNA labeling kit (Roche) was used to generate digoxigenin-labeled RNA probes for *in situ* hybridization. Constructs were linearized prior to generating probes using the listed enzyme and polymerase: *atp1a1*-antisense *Sma*I/T7 (Eid and Brandli, 2001), *lhx1*-antisense *Xho*I/T7 (Carroll and Vize, 1999; Taira et al., 1992), *hnf1*β-antisense *Sma*I/T7 (Demartis et al., 1994).

Digoxigenin-labeled *dnmbp* RNA probes were generated by first extracting DNA from stage 40 embryos as previously described (Bhattacharya et al., 2015). Regions of *dnmbp.L* and *dnmbp.S* were amplified from embryo DNA by PCR using the following primers: *dnmbp.L*- sense-Sp6 (5’ – CTAGCATTTAGGTGACACTATAGGTCAAAGGACACTCGAAACAC – 3’),

*dnmbp.L*-antisense-T7 (5’ – CTAGCTAATACGACTCACTATAGAGAAACATTCGTCTCGCGAGG – 3’), *dnmbp.S*-sense-Sp6 (5’ – CTAGCATTTAGGTGACACTATAGGTTAAAGGACACTCGAAACAC – 3’) and *dnmbp.S*-antisense-T7 (5’ – CTAGCTAATACGACTCACTATAGAGAAACGTTCGTGGAGGGTAC – 3’). PCR products were transcribed to create digoxigenin-labeled RNA probes using a DIG RNA labeling kit (Roche) and the appropriate polymerase (T7 or Sp6).

### MOs and RNA constructs

Two translation-blocking MOs were designed to target the 5’ untranslated region of *dnmbp*: Dnmbp MO 1, 5’-TCGAACCACCGATCCCACCTCCATC-3’; Dnmbp MO 2, 5’- ACCACCGACCCCACCTCCATCCTAA-3’. A standard control MO (5’-

CCTCTTACCTCAGTTACAATTTATA-3’) was used as a control for all MO experiments. MOs were ordered from Genetools. Single cell embryos were injected with 40 ng MO for Western blot analysis and 8-cell embryos were injected with 20 ng MO for phenotypic analysis.

Human *DNMBP* RNA was created by linearizing pcDNA3-HA-Tuba (Addgene plasmid 22214) DNA with *Xba*I (Salazar et al., 2003). Capped RNA for rescue and overexpression experiments was transcribed and purified from the linearized DNA using a T7 mMachine mMessage kit (Ambion). A pCS2-*β-galactosidase* construct was obtained from the McCrea laboratory for use as a control for rescue and overexpression experiments (Lyons et al., 2009a; Miller et al., 2011). *β-galactosidase* RNA was transcribed from plasmid DNA linearized with *Not*I using a Sp6 mMachine mMessage kit (Ambion).

### sgRNA design and creation

One sgRNA that was complimentary against both homeologs of *dnmbp* was designed as previously reported (DeLay et al., 2018). A sgRNA against *slc45a2* was generated for use as a control (DeLay et al., 2018). DNA templates for sgRNAs were produced by PCR, and T7 polymerase was used to transcribe sgRNA from the DNA templates as previously described (Bhattacharya et al., 2015; DeLay et al., 2018). For long-term storage, sgRNA was diluted to 1000 ng/µL and kept at -80 °C. For working stocks, sgRNA was diluted to 500 ng/µL sgRNA and stored in 5 µL aliquots of at -20 °C. Working stock aliquots were limited to five freeze-thaw cycles prior to disposal. Single cell and 8-cell embryos were injected with 1 ng Cas9 protein and 500 pg sgRNA.

### CRISPR genomic analysis

Embryos injected at the 1-cell stage with 1 ng Cas9 protein and 500 pg sgRNA were reared to stage 40. DNA was extracted from individual embryos as previously described (Bhattacharya et al., 2015), and the region surrounding the sgRNA binding site was amplified by PCR as previously described (DeLay et al., 2018). *dnmbp.L* DNA was amplified using nested PCR. The inner set of primers used were *dnmbp.L-*inner-forward (5’ – AGCTGACCCCATCTTAAAACAA – 3’) and *dnmbp.L*-reverse (5’- GTTTTTAGCTGCTTGGCTCAGT – 3’). Following the inner PCR reaction, the resulting PCR product was used to amplify *dnmbp.L* using primers *dnmbp.L*-reverse and *dnmbp.L-* outer-forward (5’ – TTCATGGCCTCTCCTACTCATT – 3’). *dnmbp.L* was sequenced using primer *dnmbp.L-*outer-forward. *dnmbp.*S DNA was amplified with primers *dnmbp.S*-forward (5’- GACCCCATAATTGAGCCATAAG – 3’) and *dnmbp.S-*reverse (5’ – CAGTGGTTTTGACGATTGTAGC – 3’) and sequenced using *dnmbp.S-*forward. TIDE was used to determine insertion and deletion frequencies in the amplified gene region (Brinkman et al., 2014).

### Microinjection

Individual blastomeres were microinjected with 10 nL of injection mix as described previously (DeLay et al., 2016). Blastomere V2 of 8-cell embryos was injected to target the kidney (Moody, 1987a). Cas9 protein (CP01; PNA Bio) and sgRNA were incubated together at room temperature for at least 5 min prior to microinjection (DeLay et al., 2018). MOs, RNA constructs and Cas9/sgRNAs were co-injected with either membrane-RFP RNA, Alexa Fluor 488 fluorescent dextran or rhodamine dextran as a tracer (Davidson et al., 2006; DeLay et al., 2018; DeLay et al., 2016).

### Immunostaining

Staged embryos (Nieuwkoop and Faber, 1994) were fixed (DeLay et al., 2016) prior to immunostaining as described previously (Lyons et al., 2009b). The lumens of the proximal kidney tubules were labeled using 3G8 antibody (1:30) and the distal and connecting kidney tubules were labeled using 4A6 antibody (1:5) (Vize et al., 1995). Additionally, proximal tubules were detected using fluorescein-labeled *Erythrina cristagalli* lectin (50 µg/mL; Vector Labs). Somites were labeled using antibody 12/101 (1:100) (Kintner and Brockes, 1984) and membrane-RFP tracer was labeled with anti-RFP antibody (1:250; MBL International PM005). Kidney, somite and membrane-RFP tracer staining were visualized using goat anti-mouse IgG Alexa 488 (1:2000; Invitrogen) and goat anti-rabbit IgG Alexa 488 and Alexa 555 (1:2000; Invitrogen).

### Imaging

Embryo phenotypes were scored and *in situ* images were taken using an Olympus SZX16 fluorescent stereomicroscope with an Olympus DP71 camera or a Leica S8 A80 stereomicroscope with a Leica MC120 HD camera. Confocal images were taken with a Zeiss LSM800 microscope. Adobe Photoshop and Illustrator CS6 were used to process images and create figures.

## RESULTS

### *Dnmbp is expressed in the developing* Xenopus *pronephros*

To assess whether Dnmbp protein is expressed during kidney development, protein lysates were collected from embryos at different developmental stages. Using a commercial antibody against Dnmbp, we found that Dnmbp protein (170 kD) is present in *Xenopus* embryos from single cell through tadpole stages by Western blot (Figure 1A). Importantly, Dnmbp protein was present from gastrula (stage 12) through tadpole (stage 38) stages when pronephric kidney specification and development occur.

**Figure 1.**
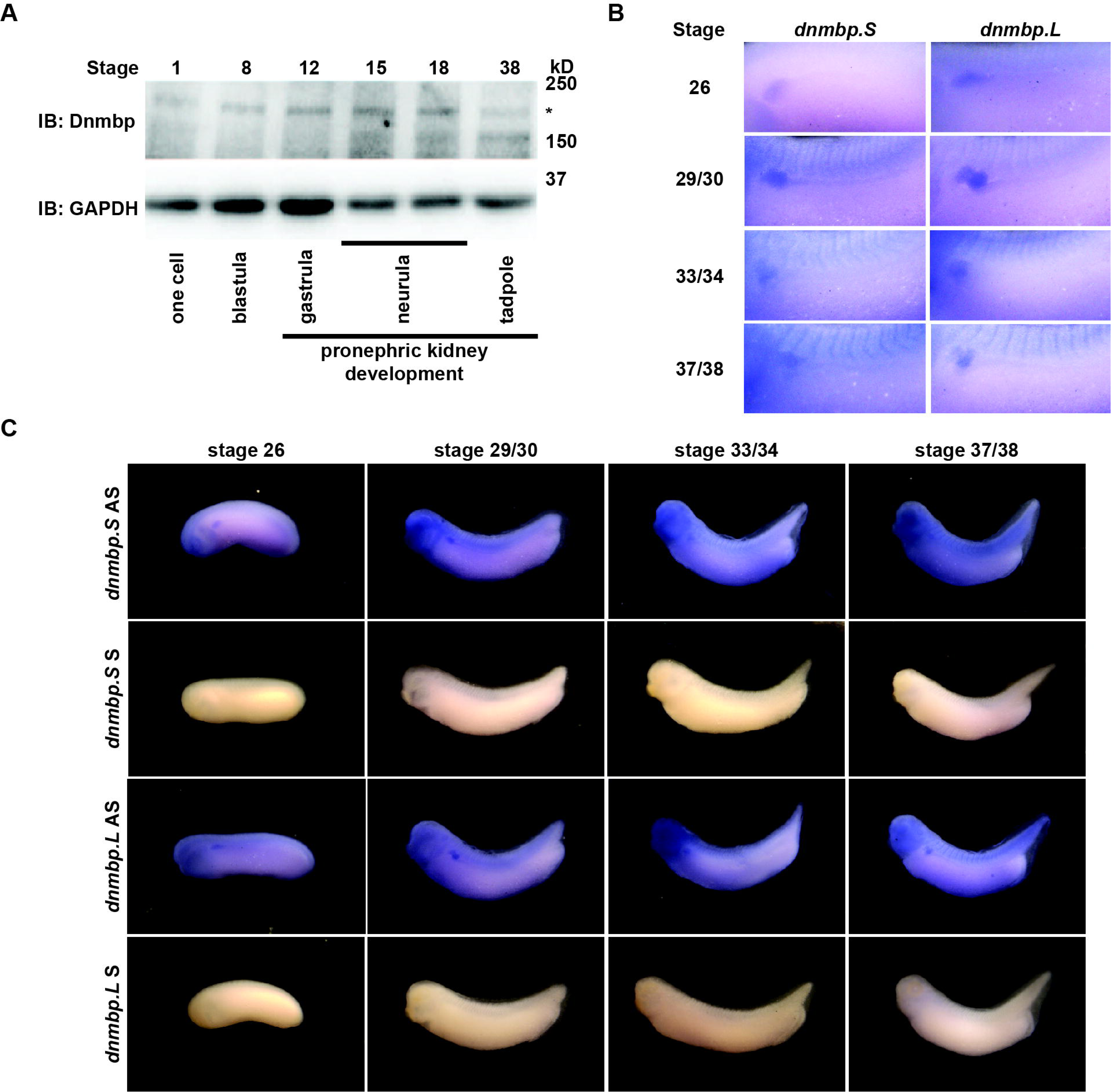
DNMBP is present throughout *Xenopus* pronephric development. A) Western blot showing that Dnmbp is present in embryos ranging from stage one to stage 38. * indicates Dnmbp. B) *In situ* hybridization using probes against *dnmbp.S* and *dnmbp.L* showing *dnmbp* expression in the developing pronephros from stage 26 through stage 37/38. C) *In situ* hybridization of both homeologs of *dnmbp*. Antisense probes (AS) labeling *dnmbp* expression in the pronephros, head structures and somites. Sense (S) probes shown as a control for non-specific probe binding were processed in parallel with the AS probes.

To determine if *dnmbp* is present in the *Xenopus* kidney throughout pronephric development, embryos ranging from stage 26 to 38 were subjected to *in situ* hybridization (Figure 1B, C). Antisense probes were created against each homeolog of *dnmbp*, and sense probes against each homeolog were used to verify that staining was specific for *dnmbp*. Starting at stage 26, both antisense probes against *dnmbp* stained the kidney tubules, with the strongest staining in the proximal tubules (Figure 1B). In addition to the kidney, the antisense *dnmbp* probes stained head structures and somites (Figure 1C). In comparison, the sense control *dnmbp* probes did not label any embryonic structures when processed in parallel to the antisense probes, indicating that the antisense *dnmbp* probe staining was specific for *dnmbp* (Figure 1C). Taken together, this indicates that *dnmbp* transcripts are present in the kidney during nephrogenesis.

### *Knockdown of Dnmbp leads to altered pronephric development in* Xenopus

To determine whether Dnmbp is necessary for the development of the *Xenopus* pronephros, we examined the expression pattern of markers of differentiated kidney tubules upon depletion of Dnmbp. Dnmbp was knocked down using two different translation-blocking MOs: Dnmbp MO 1 and Dnmbp MO 2. Both MOs were designed to target the 5’ untranslated region of *dnmbp*. Knockdown in single cell embryos was confirmed by Western blot (Figure 2A) of stage 10-12 embryos in comparison to embryos injected with a standard MO control.

**Figure 2.**
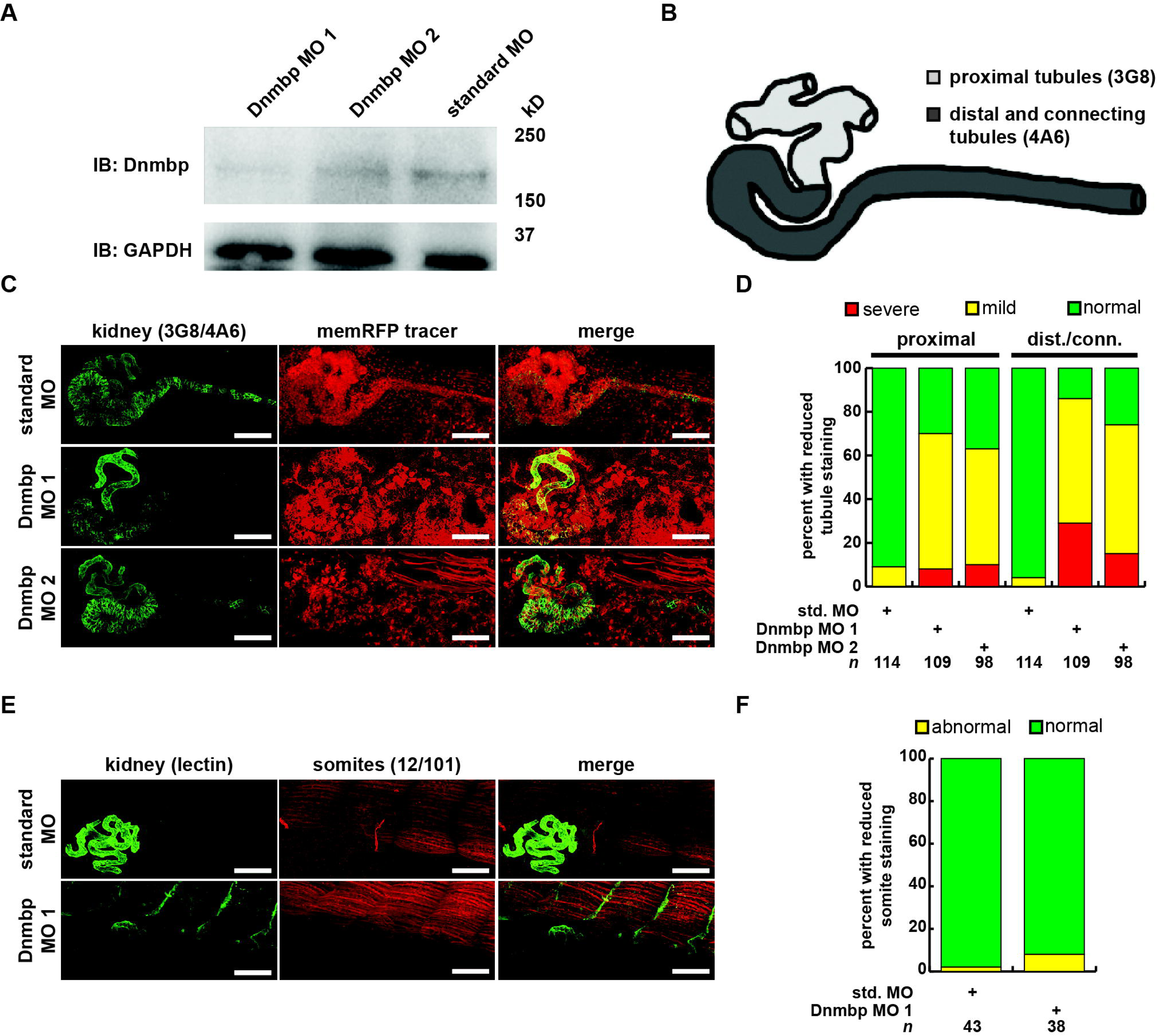
Knockdown of Dnmbp results in reduced kidney tubulogenesis, but does not alter somite development. A) Western blot showing the efficiency of two different MO targeting DNMBP. Single cell embryos were injected with 40ng of Dnmbp MO 1, Dnmbp MO 2 or standard MO. One embryo equivalent per lane was loaded on to the SDS-PAGE gel. B) Schematic of the *Xenopus* pronephros showing the proximal, distal and connecting tubule regions. C) Unilateral injection of 20ng DNMBP MO 1 and DNMBP MO 2 into blastomere V2 at the 8-cell stage leads to defects in kidney tubulogenesis in comparison to embryos injected with standard MO. Antibody 3G8 used to label the lumen of the proximal tubule, antibody 4A6 used to label the distal and connecting tubules. memRFP used as an injection tracer. White bar indicates 200µm. D) Knockdown of Dnmbp leads to reduced expression differentiated kidney tubule markers in comparison to embryos injected with standard MO. E) Unilateral injection of 20ng DNMBP MO 1 and Dnmbp MO 2 into blastomere V2 at the 8-cell stage does not cause somite defects in comparison to embryos injected with standard MO. Antibody 12/101 used to label somites, lectin used to label the proximal tubule lumen. Images are stitched from 6 tiles. White bar is 200µm. F) Knockdown of Dnmbp does not lead to reduced somite development compared to embryos injected with standard MO. *n* = number of embryos across 3 replications.

Pronephric tubule development was assessed upon Dnmbp knockdown using 3G8 and 4A6 antibodies (Vize et al., 1995), which label the differentiated proximal tubules versus the distal and connecting tubules, respectively (Figure 2B). Knockdown phenotypes of embryos injected in the left V2 blastomere at the 8-cell stage were assessed using a previously described scoring system by comparing the tubules on the MO-injected side of the embryo to the tubules on the uninjected side (DeLay et al., 2018). Phenotypes were scored as “normal” if there was no difference between the injected and uninjected side, “mild” if there was a reduction in tubule development and/or antibody staining on the injected side in comparison to the injected side or “severe” if there was little to no tubule and/or antibody staining on the injected side of the embryo.

Knockdown of Dnmbp resulted in disrupted proximal tubule development in stage 40-41 embryos that had been injected with either Dnmbp MO in comparison to standard MO-injected controls (Figure 2C and D). The proximal tubules in Dnmbp knockdown embryos had shorter branches and were less convoluted than standard MO-injected control embryos. Similarly, distal and connecting tubule development was disrupted upon Dnmbp knockdown, resulting in decreased convolution of the distal and connecting tubules of Dnmbp knockdown embryos in comparison with those of standard MO-injected controls. (Figure 2C, D). Interestingly, there was a decrease in 4A6 staining of the distal and connecting tubules even though these tubules could be visualized using the co-injected memRFP tracer (Figure 2C) indicating that although the distal and connecting tubules were present, they were less differentiated than the tubules in the standard MO-injected control embryos.

To assess whether the pronephric defects observed in Dnmbp knockdown embryos are specific, somite staining was analysed. Somites of stage 40-41 embryos were stained with 12/101 antibodies, and the lumen of the proximal tubules was stained with lectin. Embryos were injected with either standard MO or Dnmbp MO 1 at the 8-cell stage (left V2 blastomere), and somite staining on the injected side of the embryo was compared to staining on the uninjected side of the embryo. There was no difference between somite staining of standard MO- and Dnmbp MO-injected embryos (Figure 2E, F), although lectin staining indicated that there were proximal tubule defects in the Dnmbp knockdown embryos (Figure 2E). The lack of somite defects in the Dnmbp knockdown embryos indicates that the observed tubule defects are not likely to be due to secondary effects caused by somite defects.

To further assess the specificity of the Dnmbp knockdown effect, Dnmbp MO 1 was co-injected with human *DNMBP* RNA in an attempt to rescue the knockdown phenotype. *β- galactosidase* (*β-gal*) was used as a negative RNA control. Embryos were assessed at stages 40-41. Co-injection of Dnmbp MO 1 and β-*gal* RNA led to the expected decrease in proximal, distal and connecting tubule development (Figure 3A, B). Similarly, co-injection of standard MO and β-*gal* RNA did not lead to defects in tubulogenesis (Figure 3A, B). Co-injection of Dnmbp MO 1 and human *DNMBP* RNA resulted in fewer tubulogenesis defects than in Dnmbp MO 1 and β-*gal* RNA control embryos, indicating that human *DNMBP* RNA is able to rescue the phenotypic defects caused by the Dnmbp MO 1. This, combined with the lack of secondary defects shown by somite staining (Figure 2E, F), indicates that the phenotypes observed by Dnmbp knockdown are specific.

**Figure 3.**
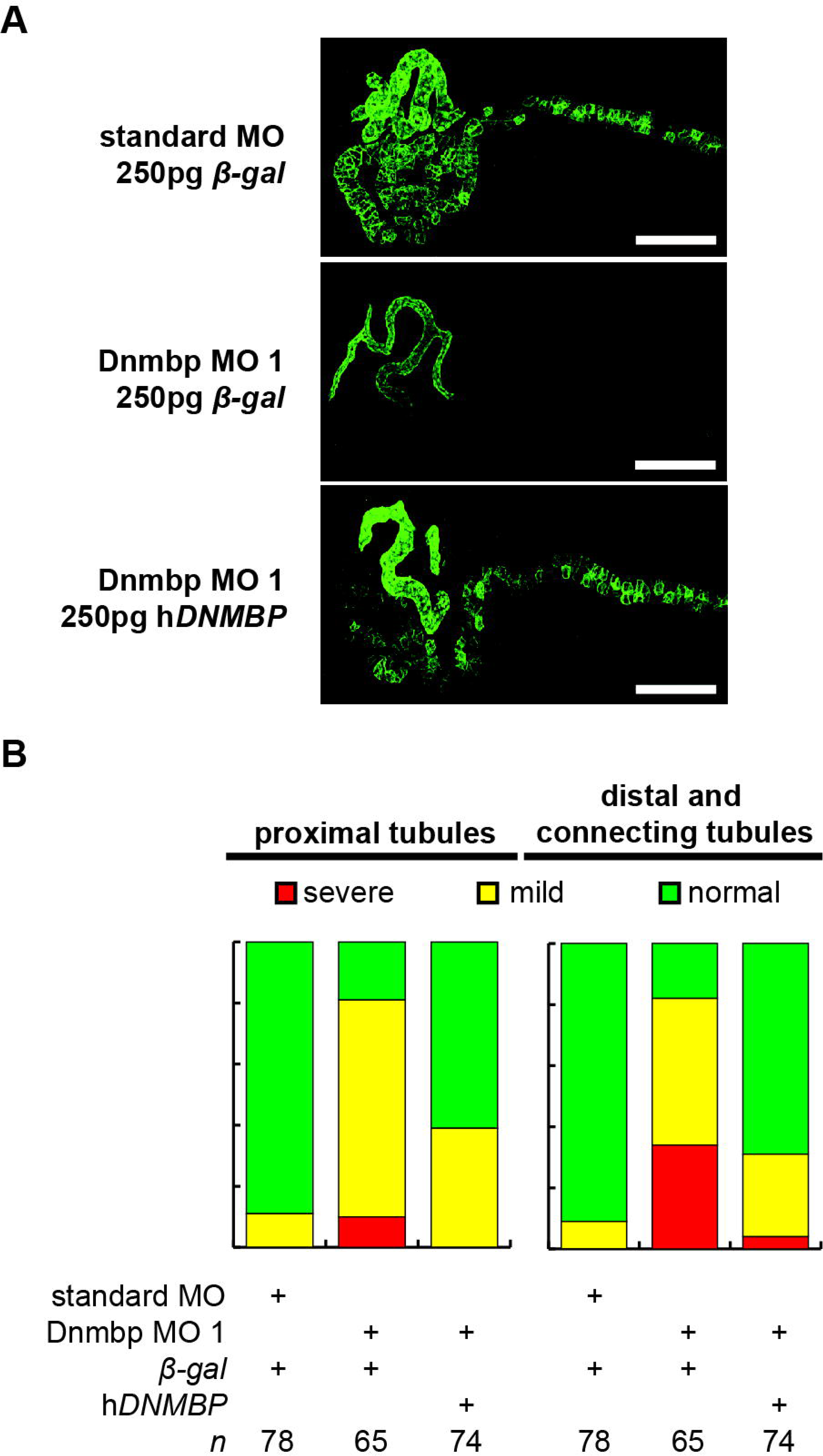
Human *DNMBP* mRNA rescues kidney defects caused by knockdown of *Xenopus* Dnmbp. A) Representative embryos showing that co-injection of *β-galactosidase* RNA with Dnmbp MO 1 does leads to kidney defects in comparison to control embryos injected with standard MO and *β-galactosidase* RNA. Co-injection of human *DNMBP* mRNA with Dnmbp MO 1 rescues the knockdown phenotype. Stage 40 embryos stained with antibody 3G8 to label the proximal tubule and antibody 4A6 to label the distal and connecting tubules. White bar is 200 µm. B) Quantitation of the rescue phenotype. *n* = number of embryos across 3 replications.

### *CRISPR* dnmbp *knockout phenocopies Dnmbp knockdown*

To further confirm that loss of *dnmbp* leads to kidney tubulogenesis defects, we designed a single sgRNA with complete complementarity to both homeologs of *dnmbp*. Embryos were injected with 500pg *dnmbp* sgRNA and 1ng Cas9 protein, and a region surrounding the sgRNA binding site was amplified by PCR. TIDE analysis of the resulting sequence indicated that the *dnmbp* sgRNA efficiently knocked out both homeologs in F0 embryos (Figure 4). Individual embryo sequence traces showed an increase in sequence trace decomposition around the expected sgRNA binding site, indicatingCRISPR editing (Figure 4A, B). Overall, CRISPR knockout of *dnmbp* resulted in 62.8% editing efficiency of the *dnmbp.L* homeolog and 61.2% editing efficiency of the *dnmbp.S* homeolog (Figure 4C, D). The most common mutation for both homeologs was an 11 base pair out of frame deletion (Figure 4C, D).

**Figure 4.**
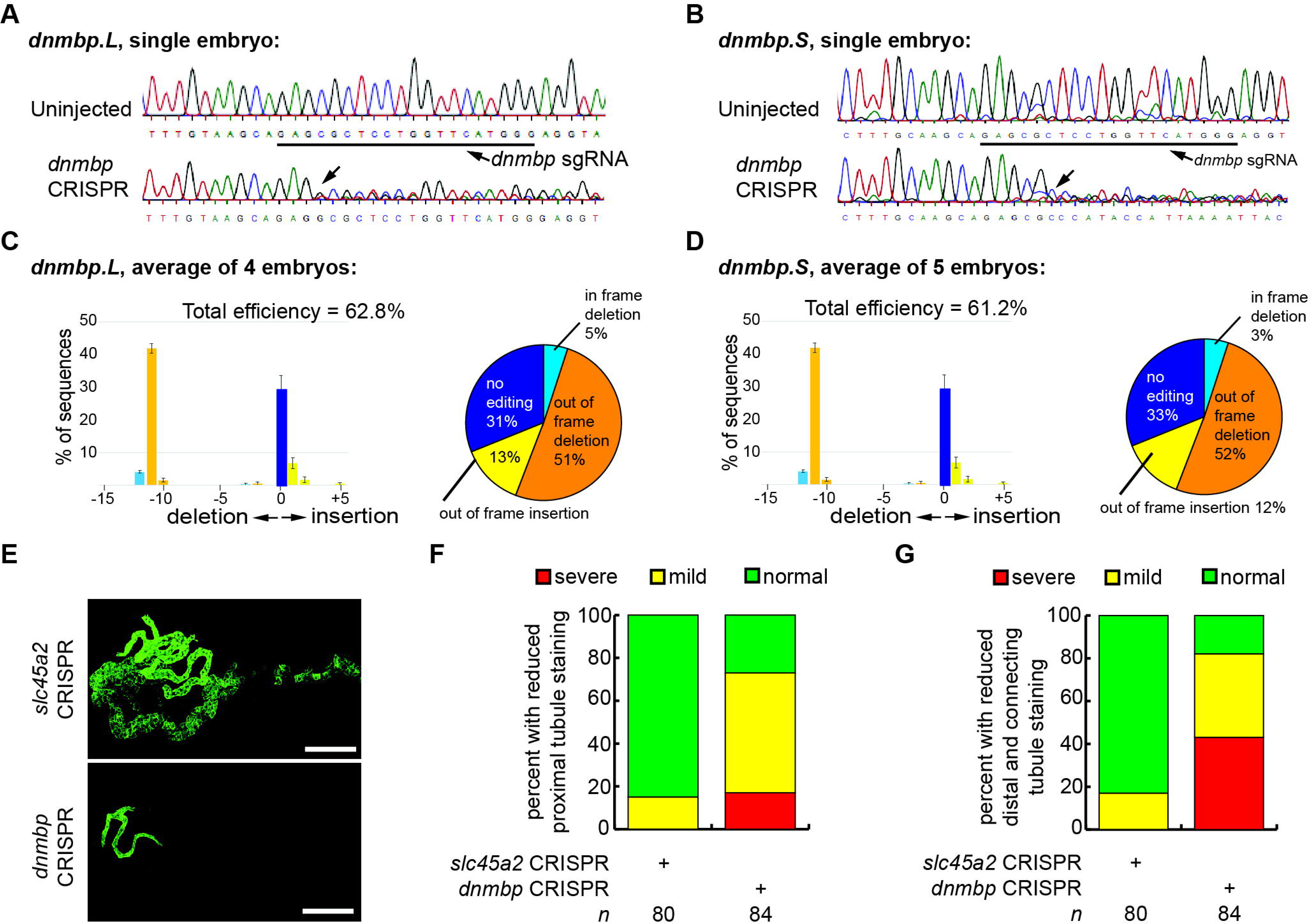
sgRNA targeting *dnmbp* efficiently edits *Xenopus* embryo DNA. Stage 40 embryos injected with *dnmbp* sgRNA and Cas9 protein at the 1-cell stage. A) Chromatogram showing CRISPR editing of *dnmbp.L* in a single embryo. The underlined sequence corresponds to the *dnmbp* sgRNA binding sequence, and the arrow indicates sequence degradation due to CRISPR. B) Chromatogram showing CRISPR editing of *dnmbp.S* in a single embryo. The underlined sequence corresponds to the *dnmbp* sgRNA binding sequence, and the arrow indicates sequence degradation due to CRISPR. C) TIDE analysis of *dnmbp.L* sequence trace degradation after the expected Cas9 cut site. * indicates p < 0.001. Percentage of *dnmbp.L* DNA sequence containing insertions and deletions. Bars indicate the mean of the percent of insertion/deletion sequences from four embryos, with the error bars representing the standard error of the mean. Results are the mean of sequencing data from four embryos. D) TIDE analysis of *dnmbp.S* sequence trace degradation after the expected Cas9 cut site. * indicates p < 0.001. Percentage of *dnmbp.S* DNA sequence containing insertions and deletions. Bars indicate the mean of the percent of insertion/deletion sequences from four embryos, with the error bars representing the standard error of the mean. Results are the mean of sequencing data from four embryos. E) Representative stage 40 embryos showing that 8-cell targeted knockout of *dnmbp* leads to disrupted kidney tubulogenesis in comparison to *slc45a2* knockout controls. Antibody 3G8 labels the proximal tubule lumen and antibody 4A6 labels cell membranes of the distal and connecting tubules. White bar is 200µm. F) Knockout of *dnmbp* reduces proximal tubule development. G) Knockout of *dnmbp* reduces distal and intermediate tubule development.

Embryos were injected at the 8-cell stage (left V2 blastomere) with 1ng Cas9 protein and either 500pg *dnmbp* sgRNA or control *slc45a2* sgRNA and reared to stage 40 to assess kidney development (DeLay et al., 2018). Proximal tubule staining of *dnmbp* knockout embryos using 3G8 antibodies indicated a reduction in proximal tubule branching and convolution in comparison to *slc45a2* knockout controls (Figure 4E, F). Distal and connecting tubule development was also disrupted in *dnmbp* knockout embryos in comparison to *slc45a2* knockout controls, with a decrease in 4A6 staining indicating less differentiated distal and connecting tubules (Figure 4E, G). The phenotype observed in *dnmbp* knockout embryos was similar to that seen in MO knockdown embryos.

### Dnmbp overexpression results in altered pronephric tubulogenesis

Knockdown and knockout of Dnmbp leads to disrupted pronephric development, so studies were carried out to determine whether Dnmbp overexpression also leads to tubulogenesis defects. Human *DNMBP* RNA was injected into single-cell embryos, which were then collected for protein lysate preparation at stage 10-12. Overexpression of human *DNMBP* RNA led to a decrease in endogenous embryo Dnmbp, with higher levels of human *DNMBP* overexpression leading to a greater decrease in endogenous Dnmbp protein (Figure 5A).

**Figure 5.**
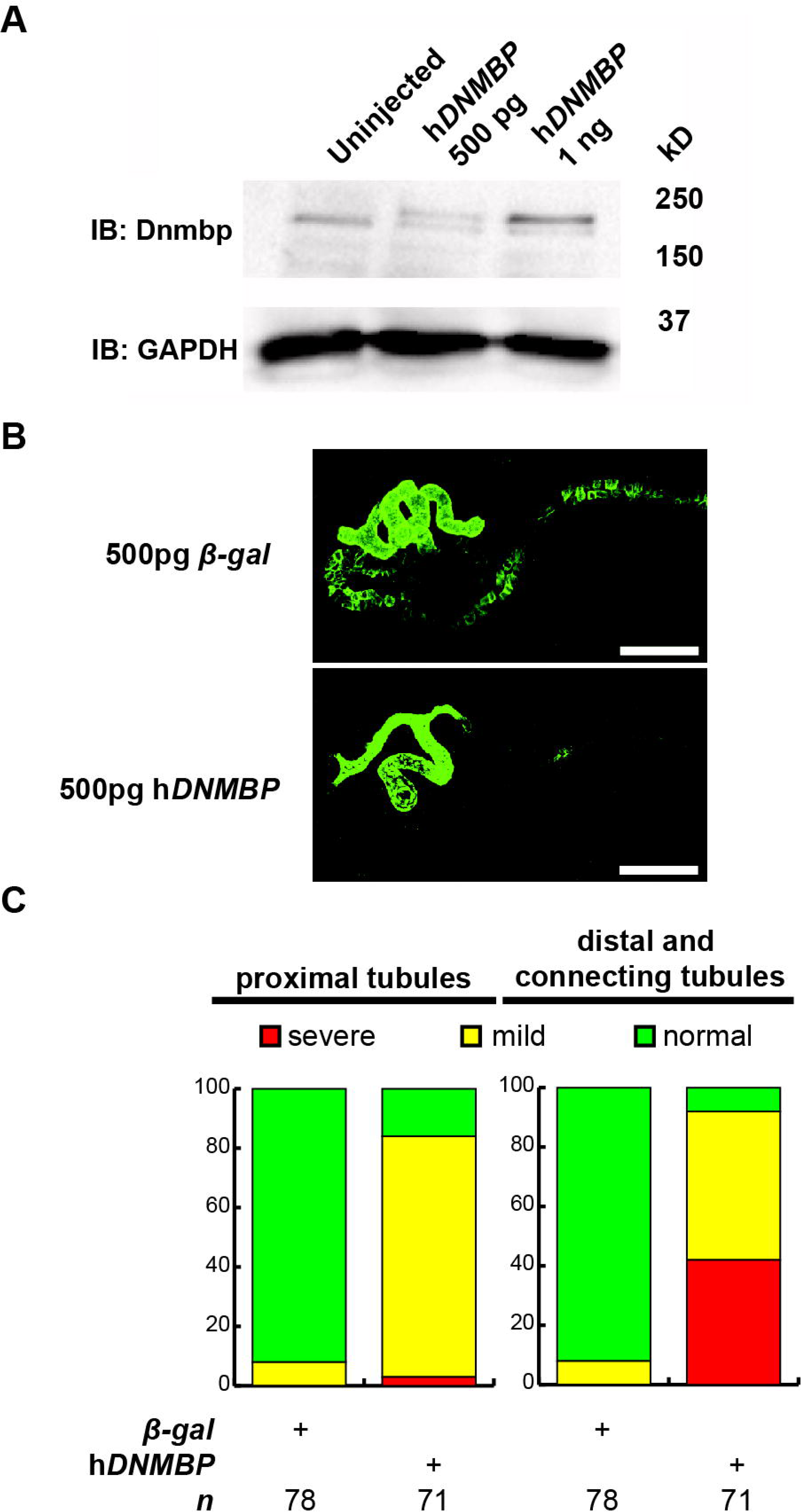
Overexpression of human *dnmbp* results in kidney tubule defects. A) Western blot showing expression of both endogenous (*Xenopus*) Dnmbp (lower band) and exogenous (human) HA::Dnmbp (upper band). Embryos injected at the one-cell stage and assayed at stage 10-12. One embryo equivalent loaded per lane. B) Representative stage 40 embryos injected with either *β-galactosidase* control mRNA or human *DNMBP* mRNA. Overexpression of *dnmbp* leads to a reduction of kidney tubulogenesis in comparison to controls. Embryos injected at the 8-cell stage (blastomere V2) to target the kidney and stained with 3G8 antibodies to label the proximal tubule and 4A6 antibodies to label the distal and connecting tubules. White bar indicates 200µm. C) Quantitation of phenotypes showing that overexpression of DNMBP leads to a reduction in proximal, distal and connecting tubule staining. *n* = number of embryos across 3 replications.

Similar to depletion of Dnmbp by knockdown or knockout, overexpression of Dnmbp led to kidney tubulogenesis defects (Figure 5B). Embryos were injected with either β-*gal* RNA as a negative control or human *DNMBP* RNA at the 8-cell stage (left V2 blastomere). Proximal tubule development was assessed using antibody 3G8, and distal and connecting tubule development were assessed using antibody 4A6 in stage 40-41 embryos. Dnmbp overexpression led to less convoluted proximal tubules with shorter branches in comparison to β-*gal* RNA control embryos. Likewise, Dnmbp overexpression led to less convoluted distal and connecting tubules, as well as decreased 4A6 staining indicating that the distal and connecting tubules were less differentiated than in β-*gal* RNA controls.

### Dnmbp depletion does not alter expression of early markers of nephrogenesis

To further understand the role that Dnmbp plays in *Xenopus* kidney development, early markers of nephrogenesis were assessed by *in situ* hybridization. Embryos were injected with either Dnmbp MO 1 or standard MO at the 8-cell stage (left V2 blastomere) and allowed to develop to stage 29-30 (*lhx1*), 32-33 (*hnf1*β) or 40-41 (*atp1a1*). Marker expression on the injected side was compared to the uninjected side of the embryo. Knockdown of early markers of pronephric development, *lhx1* and *hnf1*β, did not result in reduction of marker expression (Figure 6). This indicates that loss of *dnmbp* does not alter early kidney specification and development. Although early embryos did not show nephrogenesis defects, embryos stained at stage 40-41 with *atp1a1* showed a marked decrease in proximal tubule expression (Figure 6). The kidneys of these later stage embryos were less convoluted, but did not exhibit a loss of staining as was observed using the 4A6 antibody. This further indicates that the distal and connecting tubules are present in embryos depleted of Dnmbp, but that these tubules regions are less differentiated than they are in control embryos.

**Figure 6.**
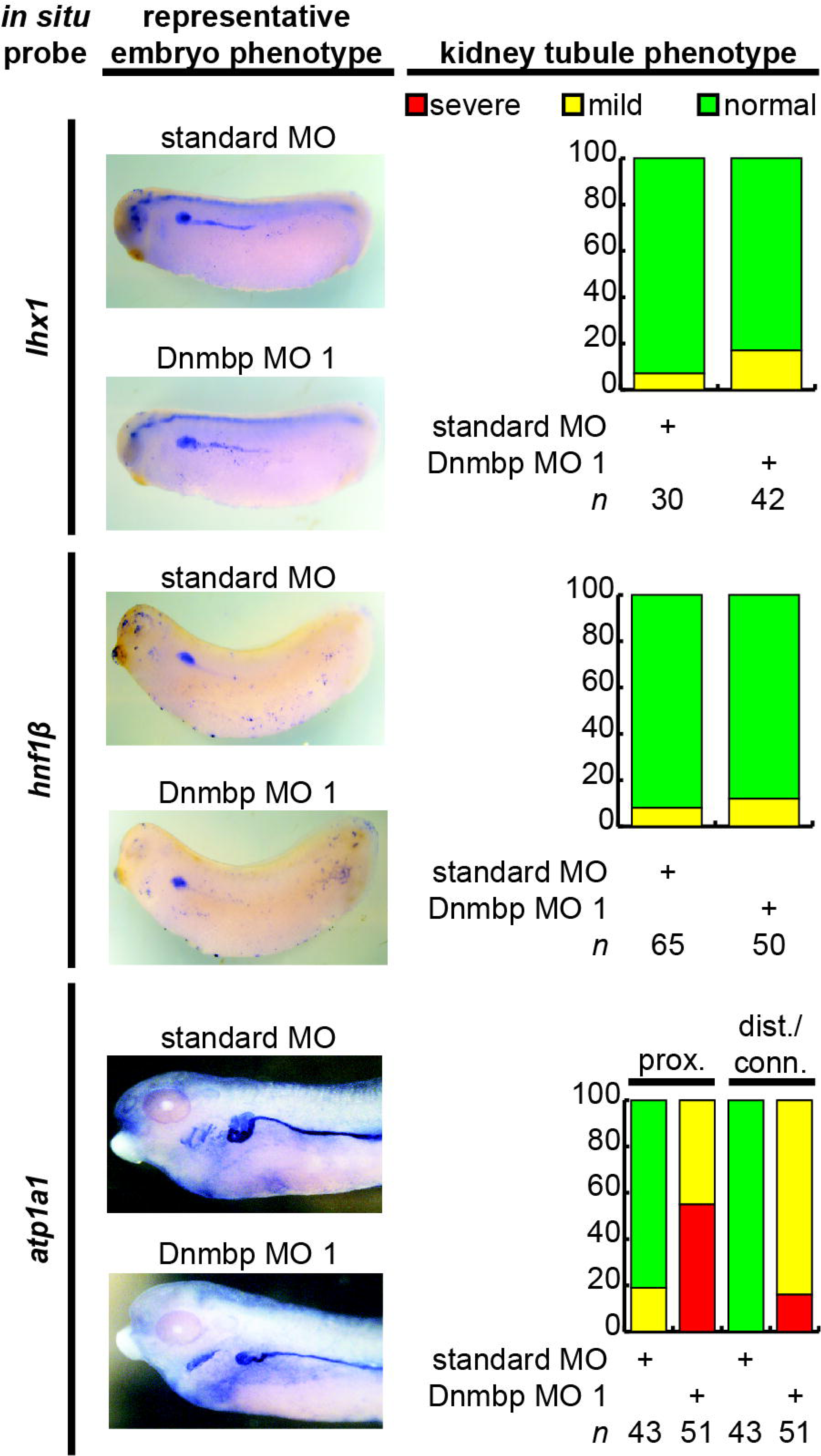
*dnmbp* knockdown does not alter expression of early markers of kidney development, however late marker expression is altered by *in situ* analysis. Embryos injected unilaterally at the 8-cell stage (blastomere V2) to target the kidney and reared to stage 29-30 for *lhx1* expression assessment, stage 32-33 for *hnf1*β expression assessment and stage 40 for *atp1a1* expression assessment. Representative embryo phenotypes shown. *n* = number of embryos across 3 replications.

## DISCUSSION

Dnmbp was first discovered in a yeast two-hybrid screen designed to identify ligands that interact with EVL, a member of the Ena/VASP family of proteins (Salazar et al., 2003). Subsequent work determined that *DNMBP* transcripts were highly expressed in human kidney tissue, in addition to other organs such as the heart, brain, lungs and liver (Salazar et al., 2003). DNMBP directly interacts with actin regulatory proteins such as N-WASP and ENA/VASP and specifically activates CDC42, thereby playing a role in the assembly of actin (Salazar et al., 2003). In the kidney, Dnmbp depletion is associated with defects in ciliogenesis and tubulogenesis (Baek et al., 2016; Zuo et al., 2011).

Here, we describe the role that Dnmbp plays in *Xenopus* pronephric development. Dnmbp protein is present in whole embryo lysates starting in single cell embryos and continuing throughout kidney development. *In situ* hybridization showed that *dnmbp* transcripts are present in the developing kidney, as well as in head structures and somites. This finding is consistent with previous work showing that *DNMBP* transcripts are present in human kidney tissue and in the zebrafish pronephros, brain and eye (Baek et al., 2016; Salazar et al., 2003). The presence of Dnmbp during *Xenopus* embryonic development, and specifically in the kidney, suggests that it plays a role in kidney development.

Knockdown of Dnmbp with two separate MOs showed that Dnmbp depletion leads to defects in *Xenopus* pronephric development. Proximal tubule branching was decreased upon *dnmbp* knockdown and the distal and connecting tubules were less convoluted than in control embryos. 3G8 and 4A6 antibodies were used to assess kidney development. Both of these antibodies label differentiated regions of the pronephric tubules, with antibody 3G8 staining the proximal tubules starting at stage 34 and 4A6 beginning to stain the distal and connecting tubules at stage 38, with complete staining by stage 41 (Vize et al., 1995). 4A6 antibody staining of the distal and connecting tubules was decreased in *dnmbp* knockdown embryos suggesting that these tubules were less differentiated than those of control embryos. These results are consistent with cell culture work that suggests that Dnmbp is necessary for tubulogenesis (Baek et al., 2016). Interestingly, previous work in zebrafish found disruption of pronephric ciliogenesis but no disruption of tubulogenesis upon *Dnmbp* MO knockdown (Baek et al., 2016). One possible explanation for this discrepancy is that the zebrafish pronephros has a less convoluted structure than the *Xenopus* pronephros (Drummond et al., 1998). Therefore, the decrease in tubule looping seen in *Xenopus* upon *dnmbp* depletion may not be apparent in the simpler zebrafish pronephros at the stages the authors examined.

The specificity of the Dnmbp MOs was confirmed by rescue experiments, where human *DNMBP* RNA was able to rescue the knockdown phenotype. Additionally, CRISPR knockout of both *dnmbp* homeologs resulted in a similar phenotype to knockdown embryos. Together, these results suggest that depletion of *dnmbp* leads to a decrease in pronephric tubulogenesis. Previous work suggests that defects in pronephric development can be secondary to defects in somite development (Mauch et al., 2000). To rule out the possibility that the kidney defects seen in knockdown embryos were the result of secondary defects due to somitogenesis defects, the somites of *dnmbp* knockdown embryos were examined. There was no difference in somite development between *dnmbp* knockdown and control embryos, indicating that the kidney defects were not likely due to larger developmental defects. This point is especially important because *dnmbp* is expressed in the somites of developing *Xenopus* embryos.

Knockdown, knockout and overexpression of *dnmbp* led to similar defects in kidney development. All three of these manipulations led to a disruption in proximal tubule development, decreased looping of the distal and connecting tubules and less differentiation of the distal and connecting tubules. These results suggest that perturbations in the level of Dnmbp of developing *Xenopus* embryo result in pronephric defects. In fact, we found that when human DNMBP was overexpressed in single cell *Xenopus* embryos, endogenous Dnmbp levels decreased. This suggests that the level of Dnmbp is tightly regulated in *Xenopus*.

Although Dnmbp knockdown results in pronephric tubule defects in stage 40 embryos, Dnmbp depletion did not result in a reduction of early markers of pronephric determination and patterning. This result indicates that the pronephros of Dnmbp knockdown embryos undergoes normal specification, and the pronephric defects seen in later embryos are due to changes in pronephric differentiation. This is supported by our *in situ* hybridization results in stage 40 embryos, where a probe for *atp1a1* completely stained the distal and connecting tubules of Dnmbp knockdown embryos, indicating that the distal and connecting tubules are indeed present. However, a loss of 4A6 staining of the distal and connecting tubules of stage 40 Dnmbp knockdown embryos indicates that the distal and connecting tubules are not completely differentiated. In conclusion, we demonstrate for the first time that Dnmbp is essential for normal vertebrate tetrapod kidney development. Our findings suggest that depletion of Dnmbp does not affect pronephric specification, but instead alters differentiation of the developing kidney tubules.

## AUTHOR CONTRIBUTIONS

B.D.D. performed microinjections, *in situ* hybridization, Western blots, designed Dnmbp sgRNA and *in situ* probes, conducted sequencing and TIDE analysis and wrote the article. T.A.B. performed initial overexpression Western blot and phenotypic experiments. R.K.M. conceived of the project, designed Tuba MOs and oversaw the experiments and manuscript preparation. All authors edited the article and approved of the final version.

## ACKNOWLEDGEMENTS

We would like to thank M.E. Corkins for performing microinjections for *in situ* experiments and critical reading of the manuscript. We are grateful for the advice and suggestions provided by members of the laboratories of R.K. Miller and P.D. McCrea, as well as M. Kloc and R.R. Behringer. We thank the University of Texas Health Science Center Office of the Executive Vice President and Chief Academic Officer and the Department of Pediatrics Microscopy Core for funding and maintaining the Zeiss LSM800 confocal microscope that was used for these studies. Additionally, we are grateful for the support of the animal care technicians and veterinarians, in particular J.C. Whitney and T.H. Gomez, who maintain our *Xenopus* colony.

